# Evaluation of *EN2* gene as a potential biomarker for bladder cancer

**DOI:** 10.1101/2022.06.28.497337

**Authors:** Ahmed Faris Aldoghachi, Aminuddin Baharudin, Umar Ahmad, Chan Soon Choy, Ong Ting Aik, Rosna Yunus, Azad Razack, Khatijah Yusoff, Abhi Veerakumarasivam

## Abstract

**Background:** Among the most prevalent cancers in the urinary tract is bladder cancer, a caner with a high rate of recurrence and metastasis as compared to other malignancies. To date, there have been many genes reported as potential bladder cancer biomarkers among which is the *EN2* gene, a member of the homeobox family containing transcriptional factors. Several studies suggested the overexpression of *EN2* to be involved with the development of a number of tumors such as bladder cancer. However, the process of involvement of *EN2* in the bladder tumorigenesis remains elusive.

**Methods:** RT-qPCR was carried out to determine the gene expression of 17 cell lines. The short-term silencing of *EN2* expression was then implemented in high-expressing cell lines using siRNAs. Using the scratch assay, the outcome of modulating the in *vitro EN2* expression on the bladder cancer migration was determined. Correlation between the IC50 values with the *EN2* expression was analyzed by correlating the viability of cells following the Newcastle Disease Virus infection with the fold change. Immunohistochemistry was then performed to determine the expression of the EN2 protein in the bladder cancer tissues.

**Results:** In the current study, *EN2* was differentially expressed in bladder cancer *in vitro* and upon modulating the expression of *EN2*, we found a reduction in the migratory effect of bladder cancer *in vitro*. In addition, following 24 hours post infection, a moderate correlation between *EN2* gene expression and NDV-mediated oncolysis was observed. No expression of EN2 in bladder cancer tissues suggesting the need for further studies to investigate the expression of EN2 protein in bladder cancer.

**Conclusion:** *EN2* may be a potential prognostic or diagnostic bladder cancer biomarker, however, further investigations are required to evaluate the *EN2* gene as a potential bladder cancer biomarker.

## 1.0 Introduction

Worldwide, bladder cancer is the sixth most prevailing cancer in males and the seventeenth in females with an approximately 573,000 new cases of bladder cancer in 2020 and 213,000 mortalities [1]. In the last decade, the incidence of bladder cancer cases and mortality rate have slightly increased [2]. It was reported that 90% of the bladder cancer patients were diagnosed with urothelial carcinoma, 5% with squamous cell carcinoma and 2% with adenocarcinoma [3]. About 70-85% of patients with an early bladder cancer diagnosis were found to have non-muscle invasive bladder cancer (NMIBC), whereas about 15-30% were found to have muscle invasive bladder cancer (MIBC) [4].

Due to the multifactorial nature of the disease, the pathogenesis of bladder cancer remains elusive to date. Among the confirmed pathogenetic factors in bladder cancer are the prolonged exposure to aromatic amines and smoking that was found to magnify the risk of bladder cancer by 2-4 times. Due to the poor prognosis, bladder cancer was reported to have the highest treatment cost in contrast to other malignancies per patient [5-6]. To date, there has been no identified prognostic and diagnostic biomarker for bladder cancer [7-8].

A homeobox is a 183 conserved base pair sequence that functions as a development regulator [9]. Among the members of the homeobox gene family is the *Engrailed Homeobox 2* (*EN2*) gene located on chromosome 7q36.3 and was found to be involved in translation, internalization, transcription, and secretion [10]. The overexpression of *EN2* has been linked to the development of breast, prostate, bladder, and colorectal cancer. Research conducted by Martin et al. 2005 revealed a high expression of *EN2* in breast cancer cell lines and a reduction in the rate of proliferation upon the suppression of *EN2* [11]. A study conducted by Morgan et al. (2011) on prostate cancer revealed a high *EN2* expression in the prostate cancer cell lines and tissues compared to normal prostate tissues/stroma and the presence of *EN2* in urine was highly associated with prostate cancer (sensitivity of 66% and specificity of 88.2%) [12]. Li et al. 2015 have identified a high *EN2* expression in the cell lines of bladder cancer [13]. In the same study, the knockdown of *EN2* revealed an inhibition in bladder tumor growth, proliferation rate and metastasis and a promotion in apoptosis and cell cycle arrest. In addition, bladder cancer tumors were found to express and secrete *EN2* which indicates its role as a potential bladder cancer biomarker due to its pathogenesis [14]. Recently, Li et al. 2020 found that *EN2* was upregulated in colorectal cancer tissues compared to adjacent normal tissues and the suppression of *EN2* resulted in inhibiting the proliferative and migratory potentials of SW480 cell line [15]. Taken together, the above studies suggest *EN2* as a potential biomarker for the diagnosis and prognosis of bladder cancer.

One of the emerging alternatives to bladder cancer therapeutics is through the use of oncolytic viruses such as the Newcastle disease virus (NDV). NDV is an enveloped avian virus of the family Paramyxoviridae with a single stranded RNA genome. NDV was found to have an anti-neoplastic property, killing cancer cells while sparing the normal cells. Additionally, NDV was found to strongly stimulate the immune system by inducing chemokines and type I IFN, increasing the cell adhesion molecules and Major histocompatibility complex (MHC), and promoting the adhesion of antigen presenting cells as well as lymphocytes through the expression of viral glycoproteins on the surface of infected cells [16-17]. Several studies have reported the anticancer potential of NDV in breast carcinoma, melanoma, head and neck carcinoma, gastric carcinoma, fibrosarcoma, hepatocellular carcinoma, neuroblastoma, and glioma [18-24]. The purpose of the present study was to identify the potential of *EN2* as a prognostic or diagnostic biomarker for bladder cancer and to investigate the response of *EN2* towards NDV therapy through the correlation of *EN2* expression with the IC50 of NDV.

## 2.0 Methods

### 2.1 Cell lines

Seventeen bladder cancer cell lines were purchased from the American Type Culture Collection (ATCC, Manassas, USA) and the European Collection of Authenticated Cell Cultures (ECACC, Salisbury, UK). Cell lines were grown in growth mediums recommended by ATCC and ECACC procedure by supplementing with fetal bovine serum (FBS) (Thermo Fisher Scientific, Inc., Waltham, MA, USA) and Penicillin Streptomycin (Thermo Fisher Scientific, Inc., Waltham, MA, USA). The cells were incubated at 37°C in a humidified atmosphere of 95% air and 5% CO2 incubator.

### 2.2 *EN2* expression analysis in bladder cancer cell lines

RNeasy Mini Kit (Qiagen. Hilden, Germany) was used to extract the RNA following the manufacturer’s protocol. PCRmax Lambda spectrophotometer (PCRmax, Staffordshire, UK) was used to determine the purity and concentration of the extracted RNA. miScript II RT Kit (Qiagen, Hilden, Germany) was used to convert the RNA to cDNA via a thermocycler (Eppendorf, Hamburg, Germany). The PCR conditions were as follow: 60 minutes incubation at 37 ºC and 5 minutes amplification at 95ºC. The miScript SYBR Green PCR Kit (Qiagen, Hilden, Germany) was adopted to quantify the *EN2* cDNA on a Light Cycler 480 (Roche, Basel, Switzerland). EN2, SDHA, TBP and GAPDH were proprietary primers that were designed by Qiagen (Qiagen, Hilden, Germany). The samples were then subjected to quantitative Real-Time PCR (RT-qPCR) following the conditions shown in Table 1.

**Table 1:**
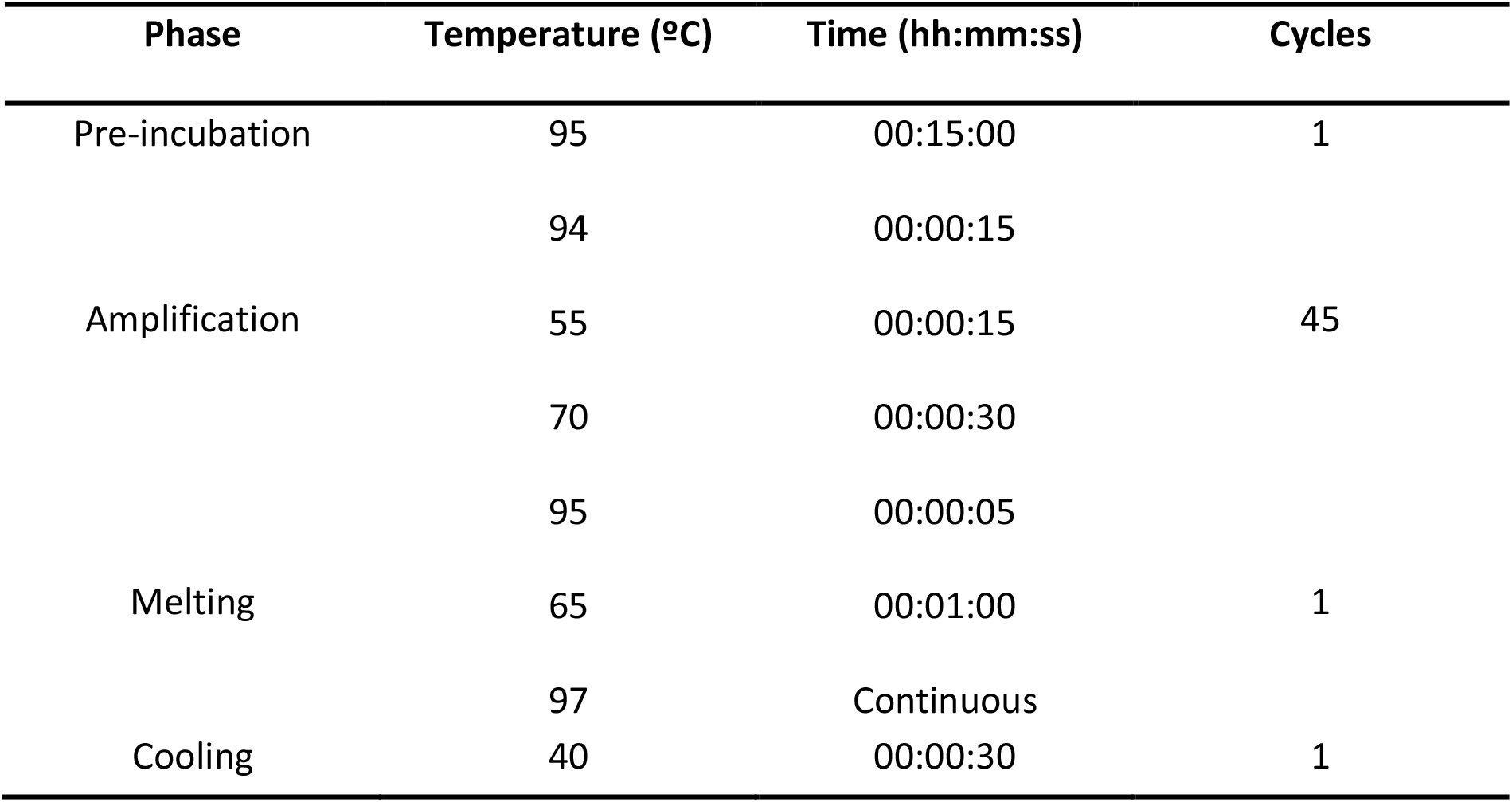
RT-qPCR temperature profile conditions.

### 2.3 *EN2* knockdown in RT112 cells

150 ×10^3^ cells/well were seeded in a 24 well-plate and replenished with fresh growth media to 600 µL in each well. Knockdown of *EN2* gene was conducted by introducing four different siRNA primers into the cells (Qiagen, Hilden, Germany) (Table 2). Four groups of cells; (1) cells transfected with siRNA, (2) cells transfected with AllStars negative control, (3) cells transfected with AllStars cell death positive control, and (4) untransfected cells were established in all the knockdown experiments. To each well, siRNA mix (30.25 µL) was added and incubated under 5% CO2 at 37 ºC. The knockdown expression of *EN2* in RT112 was studied using RT-qPCR at 24, 48 and 72 hours post-siRNA transfection.

**Table 2:**
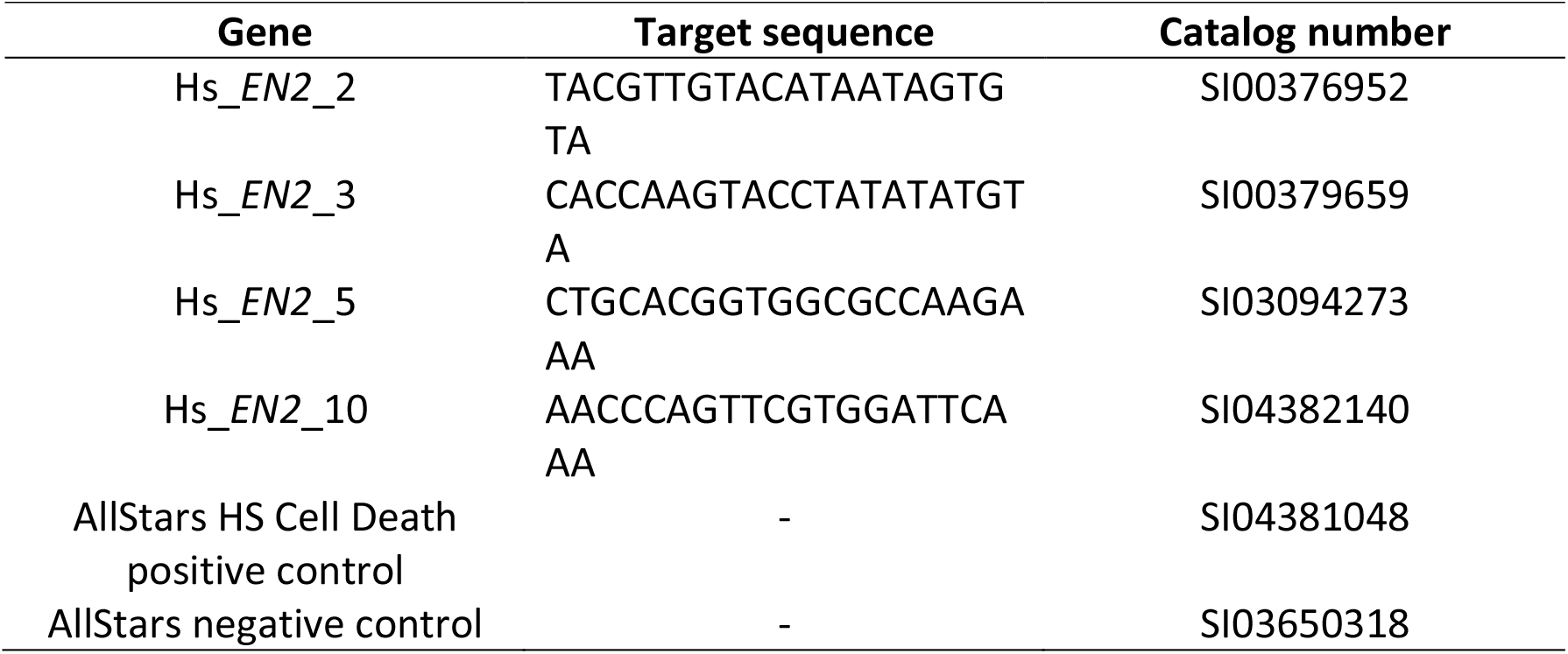
List of the siRNAs adopted in the *EN2* genes knockdown experiment along with the used controls (positive and negative).

### 2.4 Cell migration assay

The RT112 cells migratory potential was assessed using the scratch assay. A total of 150 ×10^3^ cells/well were seeded in a 24 well-plate and replenished to a total volume 600 µL in each well with fresh growth media. Prior to cell seeding, the wells were marked to enable the consistent capture of images at the same area. Cells were transfected with 31.25 µL of EN2-siRNA. The incubation of the plates then took place at 37°C with 5% CO2 until a 100% confluency was reached (72 hours). Mitomycin C (10 g/mL) (Merck, Darmstadt, Germany) was then used to treat the cells for two hours and using the tip of a 200 µL pipette, a scratch was made. Using 1X PBS, the cells were then washed, and fresh media was added into the wells. Images were then captured at three-hour intervals from the time the scratch was made until the closure of the gap for both *EN2* siRNA-transfected cells and the negative control cells (non-targeting siRNA-transfected cells). Images were then analyzed using the ImageJ software [25].

### 2.5 Correlation between EN2 expression and sensitivity towards NDV-mediated oncolysis

Using a spectrophotometer with a 440nm absorbance, WST proliferation assay was carried out to study the viability of cells infected with the AF2240 NDV strain at five different concentrations: 0, 1, 3, 5, and 7 HA units at 24, 48, 72, and 96 hours post-infection. Proliferation assay was conducted for 14 cell lines of bladder cancer namely, HTB4, UMUC5, UMUC10, UMUC1, UMUC3, UMUC13, UMUC16, TCCSUP, 5637, SCABER, RT112, J82, SW780 and HT1376. The IC50 (the concentration of virus needed to prevent the growth of cancer cells by 50%) was then determined at the four varying points of time. To correlate the gene expression and the IC50 values, the Pearson correlation was implemented, and the data was analyzed using Microsoft Excel 2010. The correlation coefficient, R^2^ was used to express the data.

### 2.6 Immunohistochemical analysis of EN2 expression in bladder cancer tissues

Formalin-fixed paraffin-embedded (FFPE) tissues from bladder cancer patients of different grades and stages were deparaffinized at 36ºC for 1 hour. Immunohistochemical analysis was carried out using polyclonal rabbit anti-EN2 ab28731 antibody (Abcam, Cambridge, United Kingdom). The slides were rehydrated in a sequence of 100, 95, 80 and 70% ethanol immersion. To retrieve the antigen, sodium citrate buffer (pH 6) (Sigma-aldrich, Missouri, USA) was then used to boil the tissue sections for 10 minutes using microwave oven and then allowed to cool at room temperature for 35 minutes. Incubation of the sections with EN2 (ab28731) antibody then took place overnight at 4ºC. Envision polymer (Envision Kit, Glostrup, Denmark) was added to the sections and incubated for 30 minutes followed by the addition of DAB chromogen (Envision Kit, Glostru, Denmark) diluted at 1:50. All sections were stained with hematoxylin and dehydrated in a sequence of 70, 80, 95 and 100% ethanol immersion and then observed under the microscope. Each IHC run included a positive and a negative control. For the positive control, a replicate of tissue slides initially identified to be positively stained for the target protein was chosen. The same criterion was used for the negative control that was incubated with 10% tris-buffered saline (TBS) overnight rather than the primary antibody. Staining intensity was scored 0-3 based on the German immunoreactive score system (IRS) [26].

### 2.7 Statistical analysis

GraphPad Prism Version 5.0.3 (GraphPad, Inc, US) was used for the statistical analysis in this study. Significant difference in the analysis of gene expression, knockdown and the migration assay of treated versus untreated groups was determined using independent t-test. The significance of EN2 protein expression in bladder cancer tissues was determined using Kruskal-Wallis test. For independent t-tests and Kruskal-Wallis test, a P <0.05 was used to measure the significance of the results. The quantitative data in the present study were expressed as mean ± standard deviation.

## 3.0 Results

### 3.1 *EN2 in vitro* gene expression

The levels of *EN2* gene expression of a number of bladder cell lines were determined. In this study, the cell lines were normalized against the HTB4 cell line (median Ct value) to determine the *EN2* expression. Results from this study demonstrated that the UMUC3 and UMUC5 were the highest *EN2*-expressing cell lines whereas UMUC10 and 5637 were significantly the lowest *EN2*-expressing cell lines (Figure 1). The fold changes of UMUC3, UMUC5, UMUC10 and 5637 relative to HTB4 were 2.60, 1.71, 0.000007 and 0.00032, respectively.

**Fig. (1).**
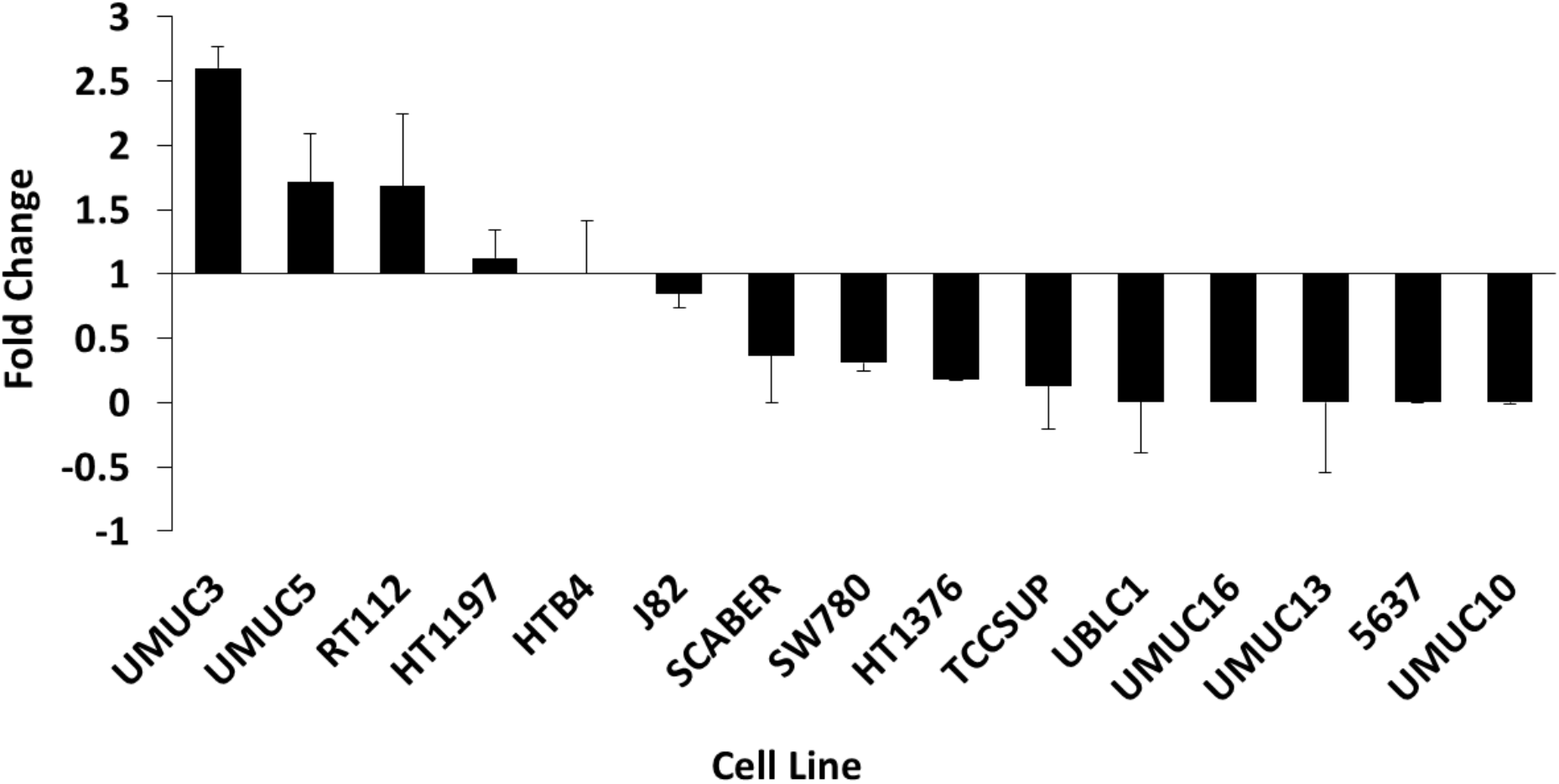
The *EN2* expression in bladder cancer cell lines. Relative expression of *EN2* gene in 16 bladder cancer cell lines measured by RT-qPCR. Cells are ordered from the highest expression to lowest expression levels. Fold change of each cell lines was calculated relative to the expression of *EN2* in HTB4 cells following the normalization with expression levels of the housekeeping genes.

### 3.2 *EN2* knockdown *in vitro*

RT-qPCR analysis was used to confirm the *EN2* expression upon transfecting RT112 cells with a combination of 4 *EN2*-targeting siRNA. The *EN2* gene levels were significantly reduced (40-48%) at 48 and 72 hours post transection in *EN2* siRNA-transfected RT112 cells relative to the non-targeting siRNA transfected negative control cells (Figure 2A). However, no significant reduction in the levels of *EN2* gene at 24 hours post transfection in *EN2* siRNA-transfected RT112 cells relative to the non-targeting siRNA transfected negative control cells.

**Fig. (2).**
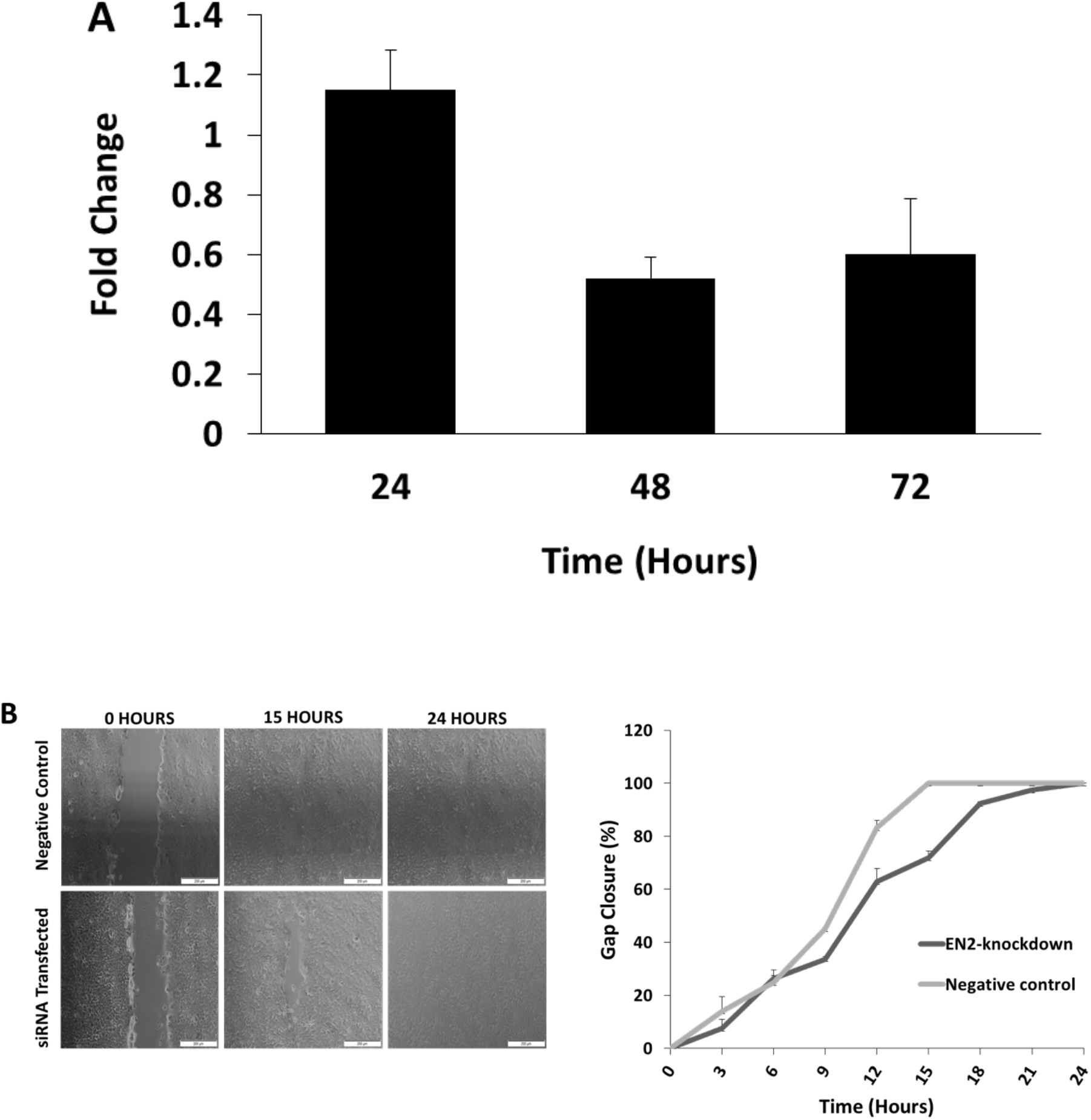
Effects of the knockdown of *EN2* on cellular migration in RT112 cells. **(A)** Relative expression of *EN2* in *EN2* siRNA-transfected cells relative to non-targeting siRNA-transfected negative control cells at 24, 48, 72 hours post-transfection. **(B)** Gap closure (%) of *EN2* siRNA-transfected RT112 cells versus non-targeting siRNA-transfected negative control in RT112 cells (200X magnification). *EN2* siRNA significantly decreased the migratory potentials of RT112 cells compared to non-targeting siRNA-transfected negative control in RT112 cells.

### 3.3 Effect of *EN2* knockdown on cell migration

Migration assay was conducted using the scratch assay to determine the effect of the knockdown of *EN2* on RT112 cell motility. A significant reduction in the gap closure rate of *EN2* siRNA-transfected cells was observed contrary to the siRNA-transfected negative control (P < 0.05). The gap for *EN2* siRNA-knockdown cells took approximately 24 hours post-scratch to close, whereas the gap of siRNA-transfected negative control cells took 15 hours (Figure 2B).

### 3.4 Correlation analysis between *EN2* expression and NDV-mediated oncolysis

Correlation analysis was carried out to identify if the *EN2* expression was correlated with the cellular sensitivity of bladder cancer towards NDV-mediated oncolysis that could serve in developing a biomarker to predict the therapeutic response. The data was expressed as correlation coefficient, R^2^ (R-squared). The obtained correlation coefficient values R^2^ were 0.4077 at 24 hours post-infection (r = 0.64; Figure 3A), 0.002 at 48 hours post-infection (r = 0.044; Figure 3B), 0.0604 at 72 hours post-infection (r = 0.24; Figure 3C), and 0.0288 at 96 hours post-infection (r = 0.16; Figure3D). The R^2^ value obtained at 24 hours indicated a moderate correlation between the IC50 values with the obtained *EN2* fold change values. On the other hand, the R^2^ values obtained at 48, 72, and 96 hours indicated a weak correlation between the IC50 values with the obtained *EN2* fold change values.

**Fig. (3).**
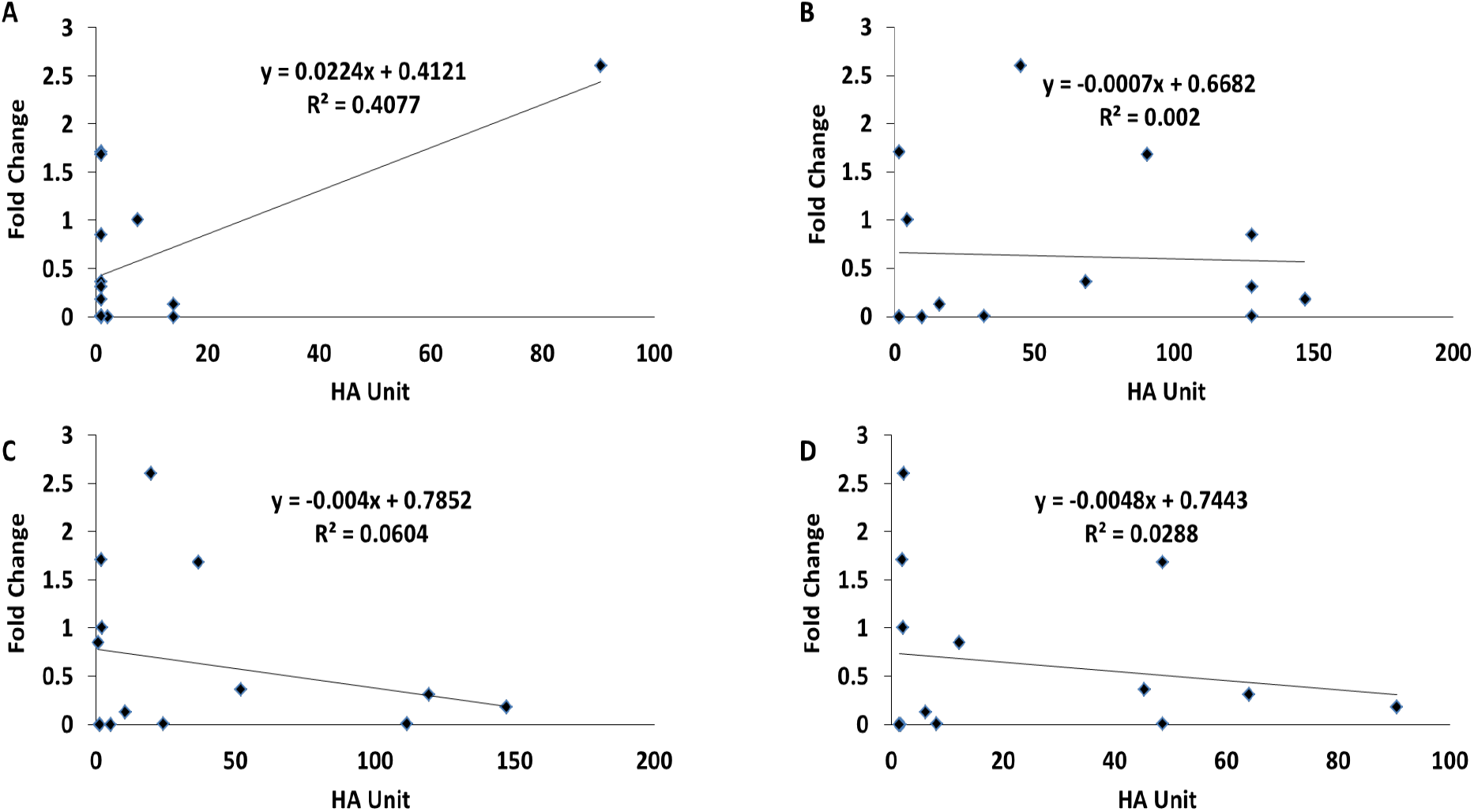
Correlation between *EN2* expression and sensitivity to NDV-mediated oncolysis. **(A)** Correlation of IC50 (HA Units) and fold change of *EN2* at 24 hours post-infection across the 14 cell lines. **(B)** Correlation of IC50 (HA Units) and fold change of *EN2* at 48 hours post-infection across 14 cell lines. **(C)** Correlation of IC50 (HA Units) and fold change of *EN2* at 72 hours post-infection across 14 cell lines. **(D)** Correlation of IC50 (HA Units) and fold change of *EN2* at 96 hours post-infection across 14 cell lines.

### 3.5 Protein expression analysis of EN2 in bladder cancer tissues

The clarity and strength of staining is affected by the concentration of the primary antibody. Therefore, optimization of the concentration of antibody to achieve specific staining with a minimal background staining was attained by using different dilutions including 1:60 and 1:200 (Fig. 4A & B). Samples were run parallel to a positive control (Fig. 4C & D). No staining for EN2 antibody was detected across all the tissue samples.

**Fig. (4).**
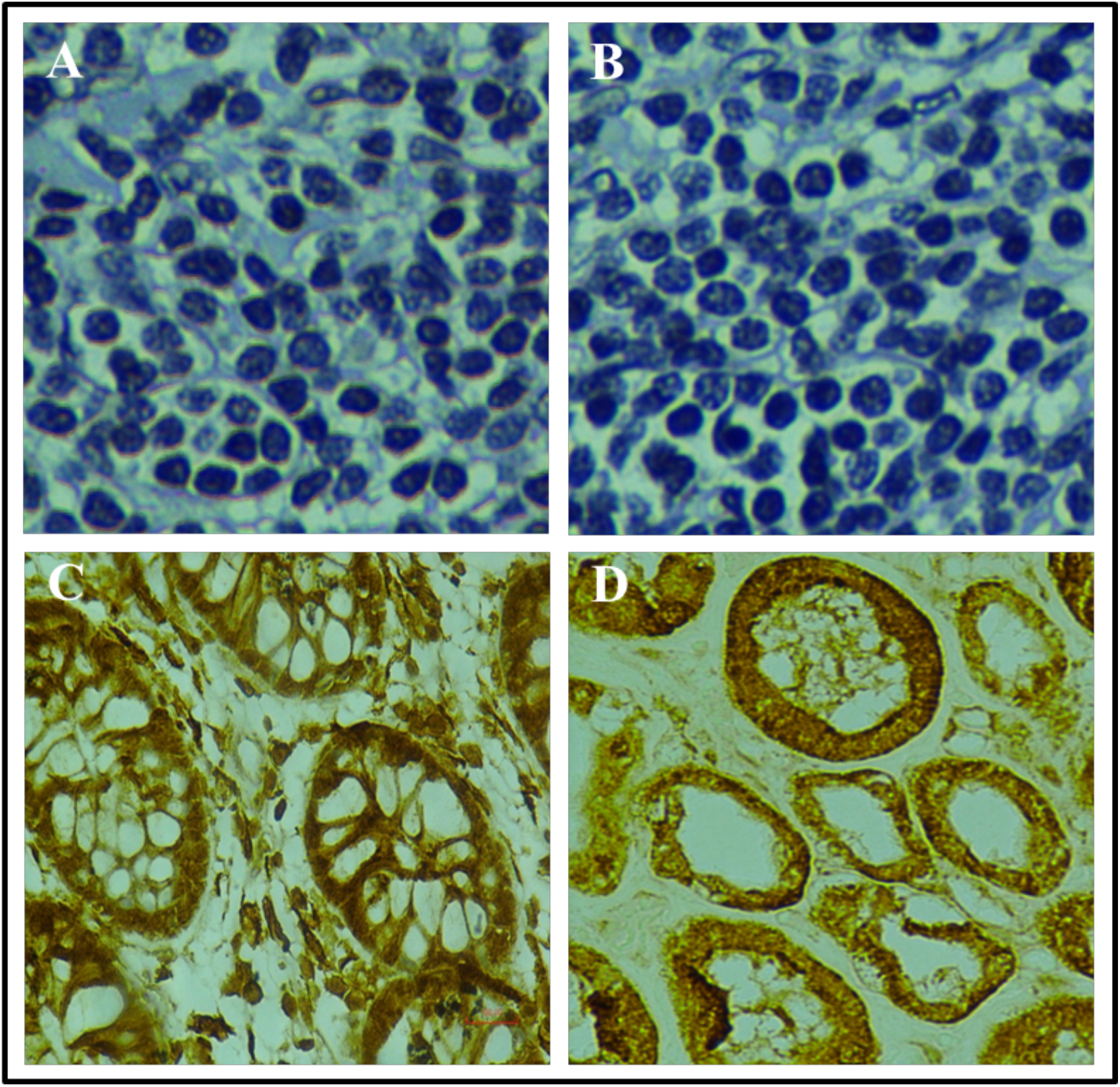
EN2 protein staining in clinical bladder cancer tissues at 400x magnification. **(A)** No protein staining with 1:60 dilution **(B)** No protein staining with 1:200 dilution **(C)** Positive control with 1:60 (Colon) **(D)** Positive control with 1:200 (Kidney).

## Discussion

*EN2*, a member of the homeobox gene family, was found to be a potential oncogene in a number of cancers [11-12]. In line with the previous studies, *EN2* has been reported to have characteristics of a potentially efficient biomarker in cancer [13,27]. *EN2* was overexpressed in human breast and prostate cancer cells, and its downregulation led to a reduction in the proliferation of prostate and breast cancer cells [11,12]. Thus, the results suggest that *EN2* expression correlates with the progression and metastasis of bladder cancer. In the current study, we have evaluated the expression of *EN2* across 17 bladder cancer cell lines and found that *EN2* was expressed in all the cancer cell lines but UMUC1 bladder cancer cells.

The obtained rank of the cell lines in Table 3 was according to the levels of *EN2* expression and represented based on histology, genetic instability, molecular subtype, and the status of classical genes mutation (*FGFR3, PIK3CA, HRAS, KRAS, NRAS, CDKN2A, TP53*) associated with bladder cancer as determined and catalogued by previous studies [28-30] (Table 1). In this study, out of the 17 used cell lines, SCaBER was the only squamous cell carcinoma which expressed *EN2* at low levels whereas the remaining cell lines were urothelial cell carcinoma. In terms of primary tumor, molecular subtype, and genetic stability, no obvious trend was observed among the high and low *EN2* expressing cell lines. As for the classical gene’s mutation, SW780 was the only cell line with no mutation in TS’P53 gene and HTB4 was the only cell line with the *HRAS* mutation. Finally, a higher frequency of the *CDKN2A* mutation variation of the *EN2* gene expression was observed among the high expressing cell lines. The tumor heterogeneity was reflected by the status of the classical gene’s mutation indicating a discrepancy in the gene expression across the various cell lines.

**Table 3.**
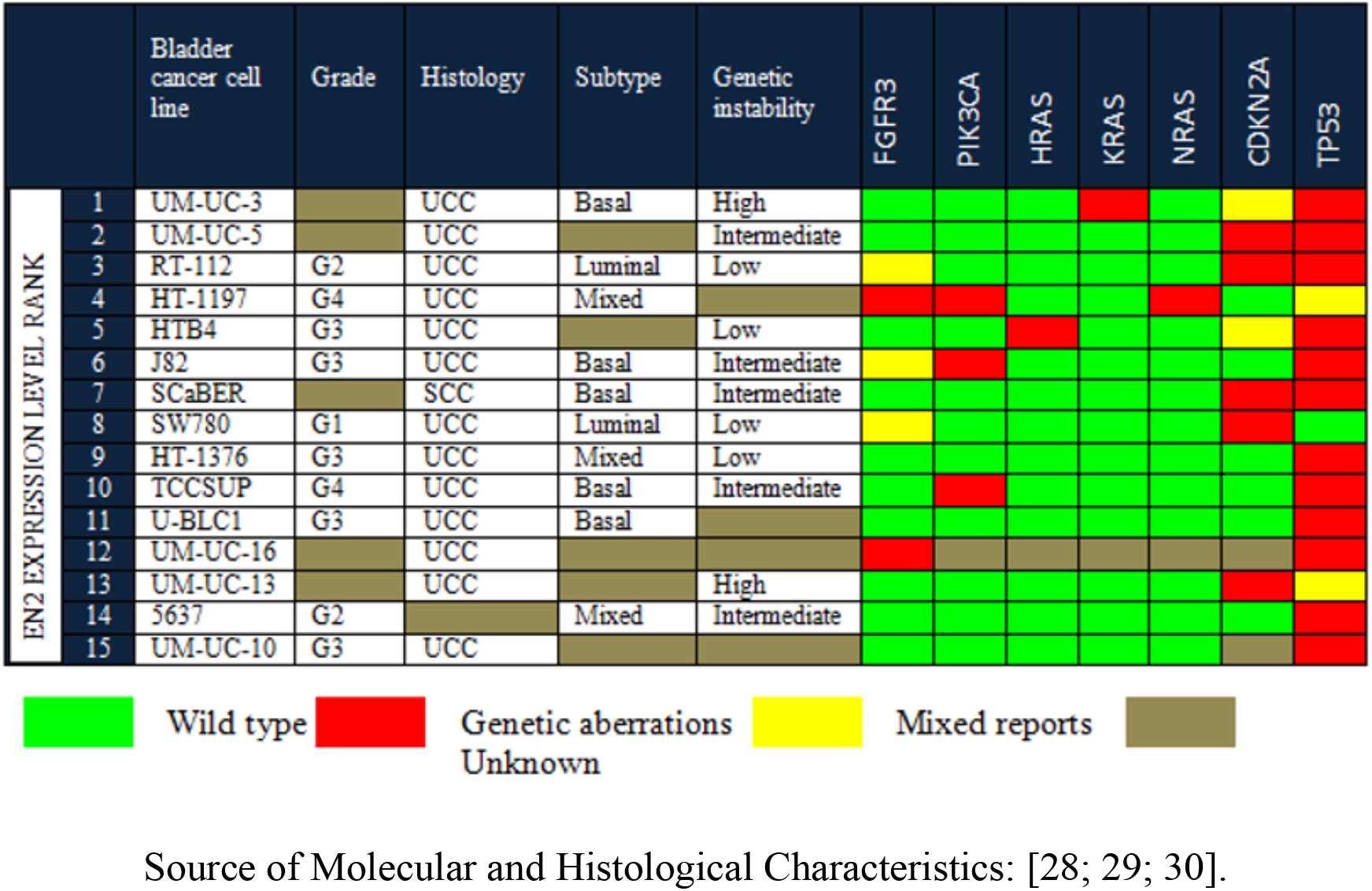
Molecular and histological characteristics of bladder cancer cell lines ranked based on the levels of *EN2* expression (Rank1 referring to the highest expression and rank 15 to lowest expression).

SiRNA induced knockdown was conducted on the RT112 bladder cancer cell line overexpressing *EN2* with the aim of clarifying the function of *EN2* in the migratory potential of bladder cancer cells. The choice of the RT112 cells instead of the UMUC3 or UMUC5 cells to conduct the knockdown was due to the knowledge that the UMUC3 and UMUC5 cells were slow growing cells. A significant knockdown of 40-48% was achieved at 48 and 72 hours post transfection. On the contrary, a higher *EN2* knockdown leading to a significant phenotypic change was achieved by Li et al. when using the siRNA sequence from Sigma (Shanghai, China) which resulted in a knockdown of 91.7% [13]. However, the discrepancy in both of the knockdown findings may not be due to siRNA alone but may as well be due to the type of cell line that might have played role in its response to the siRNA [31]. Furthermore, in the current study, the knockdown of the *EN2* expression in RT112 cells led to the decrease in the migratory potential of the RT112 cells. This result provided an insight on the possibility of *EN2* being a migratory biomarker for bladder cancer. Our findings support the role of *EN2* gene in cancer cell phenotypes as a study by Li et al. in both in vitro and in vivo experiments showed an inhibition in the bladder cancer cells invasion and proliferation following the downregulation of *EN2* via promotion of cell cycle arrest leading to apoptosis [13].

NDV AF2240 strain had been effectively tested to mediate oncolysis against breast cancer cells [24]. In our study, we found a moderate correlation between the expression values of *EN2* and the IC50 values of NDV mediated-oncolysis which was only obtained at 24 hours post infection. However, a weak or no correlation was obtained at 48, 72 and 96 hours. The moderate or weak correlation observed indicated that the *EN2* gene is not an ideal biomarker to predict the response of bladder cancer towards NDV anti-cancer therapy.

To further determine the role of *EN2* as a bladder cancer biomarker, immunohistochemistry was conducted for EN2 protein to determine the expression patterns in bladder cancer lesions including the malignant and non-malignant tissues. Our findings revealed that the EN2 protein was not expressed across all the tissues tested. Findings from our study contradicted the findings from a preceding study by Morgan et al. 2013 whereby in their study, they showed that EN2 protein was expressed at both the cytoplasmic and the nuclear level [14]. Among the possible reasons of the absence of protein expression in our findings is the long storage of tissues (2-3 years) which may have led to the antigen degradation [32]. Also, the absence of EN2 across the various grades and stages of the samples of bladder cancer may not portray an accurate expression level of EN2 in bladder cancer tissues. Other reasons may include the differences throughout the experiments that could be classified as methodological error sources and/or assessment error sources. Methodological errors may have resulted from fixation, antigen retrieval, sectioning, or antibody dilution [33]. As an example, for the removal of the tumor tissues either through biopsy or cystectomy, samples must be soaked in formalin immediately for fixation. There were 2 error sources that could have occurred in the previous step namely uneven fixation and discrepancy in the duration of the sample fixation. As for the uneven fixation, huge samples of cystectomy got a restricted contact to formalin as compared to the small biopsy chunks that possessed a higher surface area, thus better fixation due to a greater access to formalin. Regarding the differences in the duration of the sample fixation, the time variations were not recorded in Hospital Kuala Lumpur. All in all, further studies with a larger sample size from different populations must be tested to confirm the absence of EN2 protein in the tissues of bladder cancer.

## Conclusion

In our study, we have successfully showed that there were differential *EN2* expressions across different cell lines of bladder cancer. Furthermore, the knockdown of *EN2* gene reduced the migratory potential of RT112 cells. A moderate correlation between *EN2* gene expression and the IC50 of NDV-mediated oncolysis was observed at 24 hours but no correlation was observed at 48, 72 and 96 hours post-infection respectively. Also, the EN2 protein was not expressed in any of the bladder cancer tissues. It is suggested that *EN2* may potentially serve as a candidate diagnostic or prognostic biomarker for bladder cancer however, further investigations are required to clarify the patterns of EN2 protein expression in bladder cancer histopathological tissues.

## Ethical approval

The usage of human tissues in the present study was authorized by the Research and Technical Support, Ministry of Health, Malaysia (Reference number: NMRR-12-970-13096).

## Acknowledgements

This study was supported by the University of Malaysia High Impact Research (HIR) Grant Scheme: UM.C/625/1/HIR/130 and the MOSTI Flagship Fund, reference number: FP0514B0021-2(DSTIN).

## Authors’ contributions

Abhi Veerakumarasivam guided the entire project design, supervision, project administration, funding acquisition, methodology as well as writing (review and editing). Chan Soon Choy contributed in supervision, writing in terms of review and editing, and validation. Ahmed Faris Aldoghachi contributed in writing the manuscript and conducted the experimental work. Aminuddin Baharuddin, and Umar Ahmad conducted the experimental work. Khatijah Yusof provided the grant for conducting the experiments and supervision. Ong Ting Aik, Rosna Yusuf, and Azad Razack contributed in providing the ethical approvals as well as human tissue samples.

## References

[1] Sung H, Ferlay J, Siegel RL, Laversanne M, Soerjomataram I, Jemal A, Bray F. Global cancer statistics 2020: GLOBOCAN estimates of incidence and mortality worldwide for 36 cancers in 185 countries. CA: a cancer journal for clinicians. 2021 May;71(3):209–49.

[2] Bray F, Ferlay J, Soerjomataram I, Siegel RL, Torre LA, Jemal A. Global cancer statistics 2018: GLOBOCAN estimates of incidence and mortality worldwide for 36 cancers in 185 countries. CA: a cancer journal for clinicians. 2018 Nov;68(6):394–424.

[3] Kim YS, Maruvada P, Milner JA. Metabolomics in biomarker discovery: future uses for cancer prevention.

[4] Witjes JA, Compérat E, Cowan NC, et al. EAU guidelines on muscle-invasive and metastatic bladder cancer: summary of the 2013 guidelines. European urology. 2014 Apr 1;65(4):778–92.

[5] Mossanen M, Gore JL. The burden of bladder cancer care: direct and indirect costs. Current opinion in urology. 2014 Sep 1;24(5):487–91.

[6] Darwiche F, Parekh DJ, Gonzalgo ML. Biomarkers for non-muscle invasive bladder cancer: Current tests and future promise. Indian journal of urology: IJU: journal of the Urological Society of India. 2015 Oct;31(4):273.

[7] Stenzl A, Cowan NC, De Santis M, et al. Treatment of muscle-invasive and metastatic bladder cancer: update of the EAU guidelines. European urology. 2011 Jun 1;59(6):1009–18.

[8] Andreu Z, Oshiro RO, Redruello A, et al. Extracellular vesicles as a source for non-invasive biomarkers in bladder cancer progression. European Journal of Pharmaceutical Sciences. 2017 Feb 15;98:70–9.

[9] Abate-Shen C. Deregulated homeobox gene expression in cancer: cause or consequence?. Nature Reviews Cancer. 2002 Oct;2(10):777–85.

[10] Hanks M, Wurst W, Anson-Cartwright L, Auerbach AB, Joyner AL. Rescue of the En-1 mutant phenotype by replacement of En-1 with En-2. Science. 1995 Aug 4;269(5224):679–82.

[11] Martin NL, Saba-El-Leil MK, Sadekova S, Meloche S, Sauvageau G. EN2 is a candidate oncogene in human breast cancer. Oncogene. 2005 Oct;24(46):6890–901.

[12] Morgan R, Boxall A, Bhatt A, et al. Engrailed-2 (EN2): a tumor specific urinary biomarker for the early diagnosis of prostate cancer. Clinical Cancer Research. 2011 Mar 1;17(5):1090–8.

[13] Li Y, Liu H, Lai C, Su Z, Heng B, Gao S. Repression of engrailed 2 inhibits the proliferation and invasion of human bladder cancer in vitro and in vivo. Oncology reports. 2015 May 1;33(5):2319–30.

[14] Morgan R, Bryan RT, Javed S, et al. Expression of Engrailed-2 (EN2) protein in bladder cancer and its potential utility as a urinary diagnostic biomarker. European journal of cancer. 2013 Jun 1;49(9):2214–22.

[15] Li Y, Liu J, Xiao Q, Tian R, Zhou Z, Gan Y, Li Y, Shu G, Yin G. EN2 as an oncogene promotes tumor progression via regulating CCL20 in colorectal cancer. Cell death & disease. 2020 Jul 30;11(7):1–1.

[16] Washburn B, Schirrmacher V. Human tumor cell infection by Newcastle Disease Virus leads to upregulation of HLA and cell adhesion molecules and to induction of interferons, chemokines and finally apoptosis. International journal of oncology. 2002 Jul 1;21(1):85–93.

[17] Zamarin D, Palese P. Oncolytic Newcastle disease virus for cancer therapy: old challenges and new directions. Future microbiology. 2012 Mar;7(3):347–67.

[18] Ahlert T, Schirrmacher V. Isolation of a human melanoma adapted Newcastle disease virus mutant with highly selective replication patterns. Cancer research. 1990 Sep 15;50(18):5962–8.

[19] Lorence RM, Katubig BB, Reichard KW, et al. Complete regression of human fibrosarcoma xenografts after local Newcastle disease virus therapy. Cancer Research. 1994 Dec 1;54(23):6017–21.

[20] Zulkifli MM, Ibrahim R, Ali AM, et al. Newcastle diseases virus strain V4UPM displayed oncolytic ability against experimental human malignant glioma. Neurological research. 2009 Feb 1;31(1):3–10.

[21] Altomonte J, Marozin S, Schmid RM, Ebert O. Engineered newcastle disease virus as an improved oncolytic agent against hepatocellular carcinoma. Molecular Therapy. 2010 Feb 1;18(2):275–84.

[22] Li P, Chen CH, Li S, et al. Therapeutic effects of a fusogenic newcastle disease virus in treating head and neck cancer. Head & neck. 2011 Oct;33(10):1394–9.

[23] Song KY, Wong J, Gonzalez L, Sheng G, Zamarin D, Fong Y. Antitumor efficacy of viral therapy using genetically engineered Newcastle disease virus [NDV (F3aa)-GFP] for peritoneally disseminated gastric cancer. Journal of Molecular Medicine. 2010 Jun 1;88(6):589–96.

[24] Ahmad U, Ahmed I, Keong YY, Abd Manan N, Othman F. Inhibitory and apoptosis-inducing effects of Newcastle disease virus strain AF2240 on mammary carcinoma cell line. BioMed research international. 2015;2015.

[25] Schneider CA, Rasband WS, Eliceiri KW. NIH Image to ImageJ: 25 years of image analysis. Nature methods. 2012 Jul;9(7):671–5.

[26] Remmele W. Recommendation for uniform definition of an immunoreactive score (IRS) for immunohistochemical estrogen receptor detection (ER-ICA) in breast cancer tissue. Pathologe. 1987;8:138–40.

[27] Killick E, Morgan R, Launchbury F, et al. Role of Engrailed-2 (EN2) as a prostate cancer detection biomarker in genetically high risk men. Scientific reports. 2013 Jun 24;3:2059.

[28] Earl J, Rico D, Carrillo-de-Santa-Pau E, et al. The UBC-40 Urothelial Bladder Cancer cell line index: a genomic resource for functional studies. BMC genomics. 2015 Dec 1;16(1):403.

[29] Nickerson ML, Witte N, Im KM, et al. Molecular analysis of urothelial cancer cell lines for modeling tumor biology and drug response. Oncogene. 2017 Jan;36(1):35–46.

[30] Zuiverloon T, De Jong FC, Costello JC, Theodorescu D. Systematic review: Characteristics and preclinical uses of bladder cancer cell lines. Bladder cancer. 2018 Jan 1;4(2):169–83.

[31] McManus MT, Sharp PA. Gene silencing in mammals by small interfering RNAs. Nature reviews genetics. 2002 Oct;3(10):737–47.

[32] Blind C, Koepenik A, Pacyna-Gengelbach M, et al. Antigenicity testing by immunohistochemistry after tissue oxidation. Journal of clinical pathology. 2008 Jan 1;61(1):79–83.

[33] Walker RA. Quantification of immunohistochemistry—issues concerning methods, utility and semiquantitative assessment I. Histopathology. 2006 Oct;49(4):406–10.

